# Cohesin sumoylation is required for repression of subtelomeric gene expression in *Saccharomyces cerevisiae*

**DOI:** 10.1101/2025.05.26.656218

**Authors:** Deepash Kothiwal, Shalu Joseph, Shikha Laloraya

**Affiliations:** Department of Biochemistry, Indian Institute of Science, Bangalore, INDIA 560012

**Keywords:** Cohesin, gene expression, silencing, sumoylation

## Abstract

Cohesin is an evolutionary conserved protein complex first described for its role in sister chromatid cohesion, that impacts several chromosomal processes. The functions of budding yeast cohesin in chromosome segregation, replication and repair are regulated by its post-translational processing and modifications. Cohesin associates with centromeres, pericentric regions and discrete sites along chromosome arms till the end. Near chromosome ends, telomeres exist in a heterochromatin-like configuration and exert SIR-complex mediated telomere position effect resulting in repression of sub-telomeric gene transcription. Previously, we reported that cohesin has a SIR-independent role in subtelomeric gene silencing (Kothiwal and Laloraya, 2019). Here, we investigated the requirement of cohesin sumoylation in subtelomeric repression. We created a sumoylation-deficient cohesin complex by fusing the catalytic domain of a SUMO protease, *ULP1,* to the C-terminus of Mcd1/Scc1 (Mcd1-UD), a cohesin subunit. We show that cohesin sumoylation is required for repression of sub-telomeric genes. In agreement with our earlier observations of a SIR-independent role of cohesin in telomere silencing, SIR-proteins remained bound to a de-repressed subtelomeric gene in *MCD1-UD* and its expression further increased upon deletion of *SIR2*. Interestingly, we did not observe a cohesion defect in this mutant suggesting that sister chromatid cohesion and regulation of sub-telomere gene silencing are separable functions of cohesin. Telomere tethering to the nuclear envelope and telomere compaction are defective in *MCD1-UD,* indicating that sumoylation contributes to cohesin’s role in subtelomeric chromosome organization. Our data establish the relevance of cohesin sumoylation in subtelomeric repression, a function independent of cohesin’s role in sister chromatid cohesion.

## Introduction

Chromatin is a dynamic structure with varying organization along the chromatin fibre. Some regions are tightly compacted into transcriptionally inactive heterochromatin domains while others form less compact and therefore more accessible, transcriptionally active euchromatin domains. This provides an excellent mechanism through which gene expression can be regulated at the level of chromatin organization. In *Saccharomyces cerevisiae*, heterochromatin is found at the silent mating loci (*HML*α and *HMRa*, or the *HM* loci), telomeres and the rDNA. Heterochromatin contains hypoacetylated and demethylated nucleosomes that are bound by the SIR (<u>S</u>ilent <u>I</u>nformation <u>R</u>egulator) complex, which consists of Sir2, Sir3 and Sir4. Sir2 is the founding member of a conserved family of NAD-dependent protein deacetylases required for silencing at all three loci (1–5). Sir3 and Sir4 are histone-binding proteins that bind with high affinity to deacetylated and demethylated nucleosomes (2,3,6). All SIR proteins (Sir1, Sir2, Sir3, Sir4) are required for silencing of HM loci and telomeres, whereas a different protein complex known as the RENT complex (Sir2, Net1, Cdc14) is essential for rDNA silencing (7).

As first described in *Drosophila* (8), heterochromatin structure at telomeres results in transcription silencing of nearby genes, a phenomenon known as telomere position effect (TPE). In *S. cerevisiae,* TPE was first described using reporter genes placed near modified telomeres either on the left arm of chromosome VII or the right arm of chromosome V (9). The effect was shown to be independent of gene identity and promoter sequence. In *S. cerevisiae,* silencing at telomeres is established by interaction and recruitment of Sir4 by telomeric repeat binding protein Rap1 and telomere end binding proteins, Yku complex (Fig 2.1) (10,11). Sir4 in turn recruits Sir2 and Sir3, that deacetylates nearby histones thereby creating high affinity binding site for the SIR complex. This cycle continues to spread silencing in the subtelomeric region until it encounters a silencing barrier. Later studies questioned this continued spreading model and using *URA3* as silencing reporter and showed that not all telomeres are silent and that silencing is discontinuous across the length of the telomere and mostly restricted to positions close to the X element (12). Moreover, despite strong enrichment of Sir-proteins at telomeric repeats and core X elements, only subsets of subtelomeric genes were repressed in SIR-dependent manner (∼6%) (13,14). Nonetheless, it has been shown that genes present within 20 kb from the chromosome end are expressed at lower level compared to rest of the genome (13,14) confirming TPE in *S. cerevisiae,* though the mechanism and proteins involved in this repression remained largely unknown.

**Figure 1.**
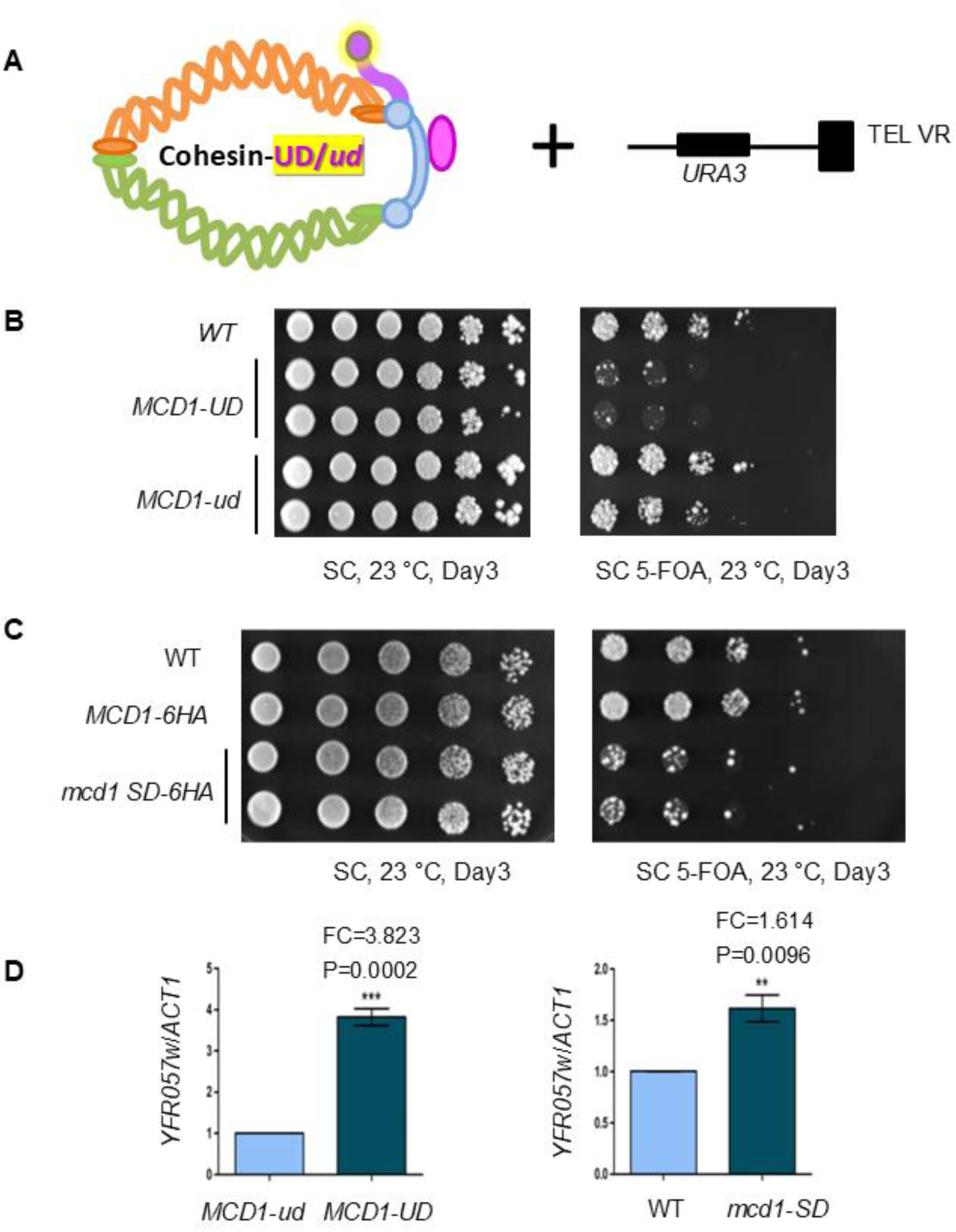
Telomere silencing defect in cohesin sumoylation defective cells. (**A**) Schematics of the cohesin-UD fusion (left) and modified telomere VR having *URA3* silencing reporter (right). The UD catalytic domain is represented by the glowing purple ball attached to the Mcd1/Scc1 subunit shown in blue. (**B**) TPE measurement in ROY783. Overnight grown log phase cultures of wild type (ROY783), *MCD1-UD* (SLY1820) and *MCD1-ud* (SLY1823) strains were serially diluted and spotted onto SC and SC + 0.1% 5-FOA plates. Plates were incubated at 23 °C for indicated time. (**C**) Spot test to analyse silencing of telomere-proximal *URA3* reporter in ROY783 at 23 °C in wild-type (ROY783), *MCD1-6HA* (SLY2743) and *mcd1 sd-6HA* (SLY2745). **(C)** Expression levels of native telomere proximal genes in *MCD1-UD* and *mcd1-SD.* RT-qPCR analysis to compare transcript levels of sub-telomeric gene, *YFR057w* in *MCD1-ud* (SLY1823) and *MCD1-UD* (SLY1820) strains (left panel), and in WT (SLY2743) and *mcd1-SD* (SLY2745) (right panel). The mean values for n≥3 experiments are plotted on the Y-axis, Error bars indicate standard error of the mean (SEM); FC∼fold-change. The statistical significance was analysed using the two-tailed unpaired t-test where * indicates P≤0.05, ** indicates P≤0.01 and *** denotes P≤0.001.

**Figure 2.**
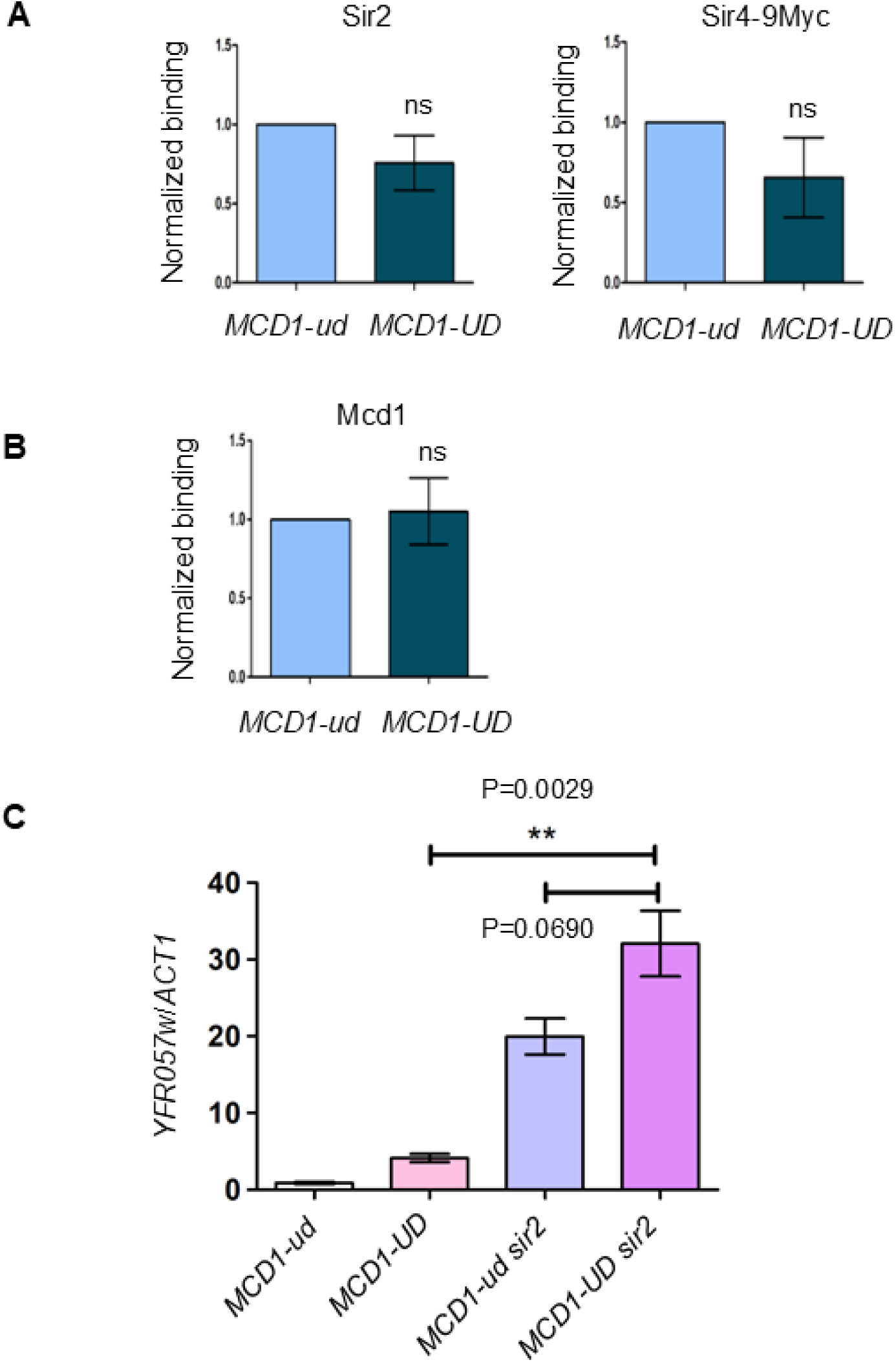
Cohesin sumoylation mediated repression of subtelomeric genes is independent of Sir proteins. (**A**) Binding of Sir proteins at *YFR057w* by ChIP-qPCR. Histograms for Sir2 and Sir4-9Myc binding are shown. In chromatin immunoprecipitations assay, the fold enrichment levels were calculated relative to *PHO5.* The mean values for n≥3 experiments are plotted on the Y-axis, Error bars indicate standard error of the mean (SEM). For simplicity wild-type values were normalized to 1. (**B**) ChIP-qPCR to analyse Mcd1 binding at 500bp from telomere VIR in *MCD1-ud* (SLY1823) and *MCD1-UD* (SLY1820) cells. **(C)** RT-qPCR analysis to measure expression level of *YFR057w* in *MCD1-ud* (SLY1823), *MCD1-UD* (SLY1820) compared to *MCD1-ud sir2* (SLY2911) and *MCD1-UD sir2* (SLY2912). Expression levels are measured relative to *ACT1.* The mean values for n=3 experiments are plotted on the Y-axis, Error bars indicate standard error of the mean (SEM). The statistical significance was analysed using the two-tailed unpaired t-test where ** indicates P≤0.01, ns = non-significant.

The cohesin complex consists of Smc1 and Smc3 along with two non Smc subunits, Mcd1/Scc1, the kleisin component of the complex, and Scc3/Irr1. Cohesin topologically embraces sister chromatids to provide cohesion (15,16). Cohesin binds to centromeres, pericentromeric regions and specific sites along the chromosome arms (17–22). In addition to its central role in sister chromatid cohesion, cohesin is also involved in various other aspects of chromosome biology such as condensation, DNA replication and repair, and regulation of gene expression (23–26). Several developmental defects (collectively known as Cohesinopathies) are associated with mutations in components of the cohesin network (27–30); and these defects could be a result of altered gene expression rather than defect in sister chromatid cohesion (31,32). Inactivation of the cohesin complex in budding yeast has also been shown to affect the expression of a small number of genes (24) and recently we showed that cohesin is required for repression of telomere proximal genes in a SIR independent manner (26).

Cohesin function is modulated by various posttranslational modifications including phosphorylation, acetylation and sumoylation. Cohesin is known to be sumoylated in both *S. cerevisiae* and humans (33–36). Wu *et al.* showed that Mcd1 sumoylation is required for sister chromatid recombination (SCR) upon DNA damage but not for mitotic cohesion in human cells (Wu et al 2012). In budding yeast, all cohesin subunits undergo sumoylation in an Mms21 dependent manner, although Siz1 and Siz2 are also required for complete sumoylation (33,36). Using an intriguing approach of fusing the catalytic Ulp domain (UD) of Ulp1 SUMO peptidase to the C-terminus of Mcd1, Almedawar et al showed that cohesin sumoylation is required for its function in providing sister chromatid cohesion. At the same time McAleenan et al also created a sumoylation-deficient version of Mcd1 by mutating 11 lysine residues to arginine. They showed that Mcd1 sumoylation is required, independent of Chk1 mediated phosphorylation, for damage induced cohesion, both at the site of a break and genome wide (33).

Having established a role of cohesin in subtelomeric gene silencing (26), we explored the role of its sumoylation in repression of telomere proximal genes. Consistent with our observation with temperature sensitive mutants of cohesin, we found that cohesin sumoylation is required for complete repression of telomere proximal genes independent of Sir proteins. Interestingly we found that sister chromatid cohesion and transcriptional repression are separable function of cohesin. Furthermore. we show that reduction in sub-telomere silencing is accompanied by telomere organization defects in this mutant.

## Results

### Fusion of Ulp domain (UD) to the kleisin subunit (Mcd1/Scc1) of the cohesin complex results in de-sumoylation of cohesin and sensitivity towards DNA damaging agents

All the cohesin subunits in *S. cerevisiae*, Smc1, Smc3, Mcd1/Scc1 and Scc3, are reported to be sumoylated (36). The budding yeast genome codes for two SUMO proteases, Ulp1 and Ulp2. In order to understand the significance of sumoylation of the cohesin complex, the catalytic domain (UD) of Ulp1 was fused to the C-terminus of Mcd1 with 3HA as a linker in-between (referred to as Mcd1-UD) (Fig S1A), using a strategy similar to Almedawar *et al*.

As the kleisin component has access to both SMC and non-SMC proteins, a Ulp domain attached to its C-terminus should be able to de-sumoylate the whole cohesin complex. As a control we created a similar fusion protein (Mcd1-UD^FA,CS^), where the Ulp domain was inactivated by mutating residues (F474A, C580S) involved in binding to SUMO and peptidase activity (Fig S1A). The steady state level of Mcd1 was unaltered upon fusion with UD, compared to Mcd1-3HA or Mcd1-UD^FA,CS^ (Fig S1B). We also tested the sumoylation status of the cohesin complex in Mcd1-UD using a strain expressing SUMO (Smt3) with a 6XHis-Flag tag at the N-terminus. All the sumoylated proteins in the cell were pulled-down using Ni-NTA resin and sumoylation of the test protein was checked by western blot analysis using anti-HA antibody in case of Mcd1 or anti-Myc antibody for Smc1 and Smc3 that were myc tagged (Fig S2A and S2B). Consistent with published results (36), fusion of the Ulp domain (UD) to Mcd1 diminished the sumoylation of all three tested subunits (Mcd1, Smc1, Smc3) of the cohesin complex compared to Mcd1-UD ^FA,CS^ (hereafter referred as *MCD1-ud)* (Fig S2A and S2B), suggesting that Mcd1-UD can indeed de-sumoylate the whole cohesin complex.

We also tested the growth phenotype of *MCD1-UD,* relative to *MCD1-ud* or wild-type cells by spot assay at various temperatures and in the presence of DNA damaging agents (Fig S3). The strain harboring the *MCD1-UD* allele showed a growth defect at 23 °C but grew comparable to *MCD1-ud* or wild-type cells at 30 °C or 37 °C (Fig 3.4). Consistent with other SUMOylation-defective cohesin mutants (33,35), *MCD1-UD* cells were sensitive towards HU and MMS (Fig S3), suggesting a role of cohesin sumoylation upon replication stress and DNA damage. Under all the conditions tested, *MCD1-ud* (catalytically inactive fusion) grew as well as wild-type, therefore in all our further studies, *MCD1-ud* was used as a wild type control.

**Figure 3.**
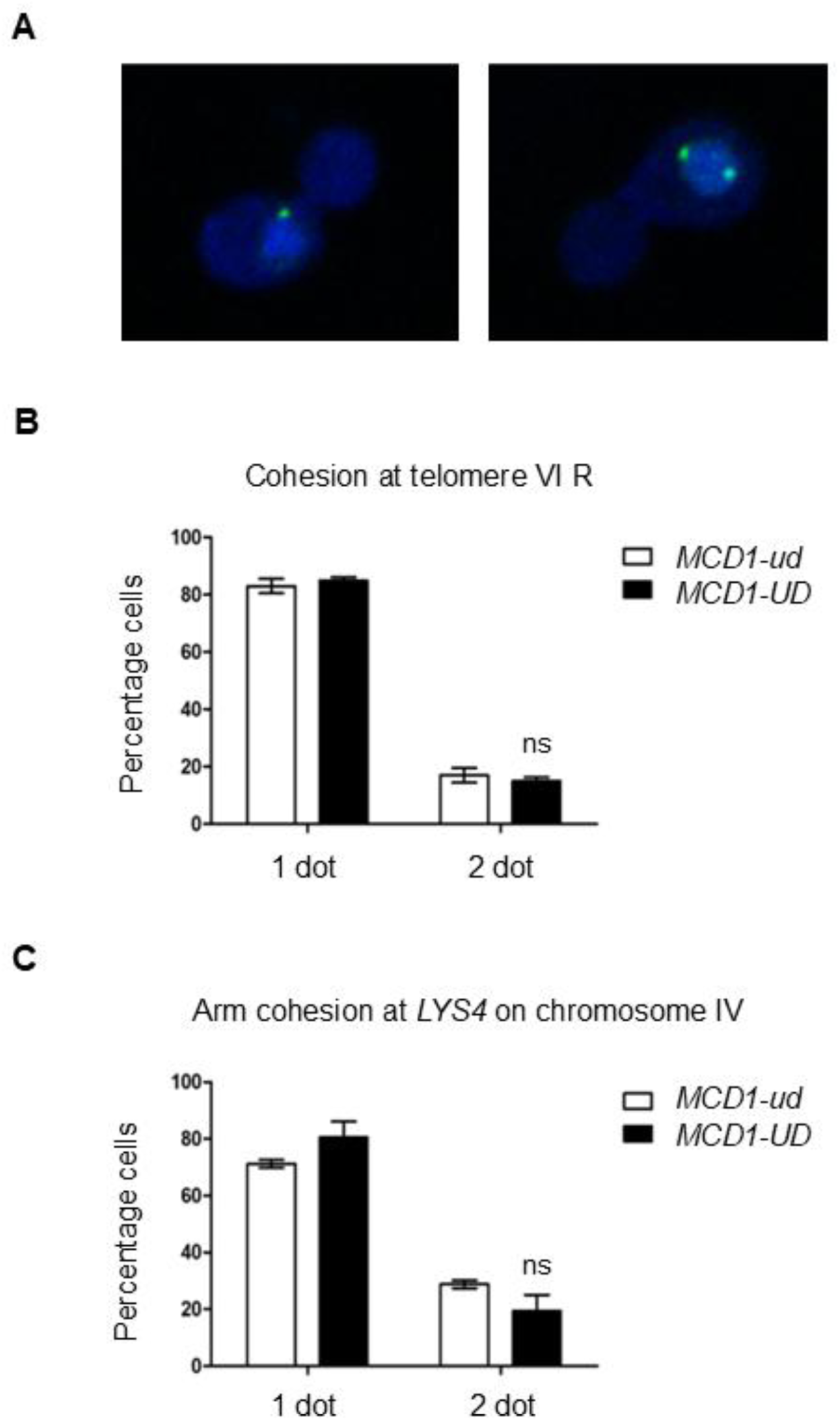
Analysis of sister chromatid cohesion in *MCD-UD* cells at 23 °C. **(A)** A merged image of DAPI and GFP showing cohesed (left) and uncohesed (right) configuration. **(B)** Analysis of cohesion at a telomere proximal site in *MCD1-ud* (SLY2549) and *MCD1-UD* (SLY2551) cells carrying a CFP chromosomal tag near telomere VIR. **(C)** Analysis of sister chromatid cohesion at *LYS4* in *MCD1-ud* (SLY2557) and *MCD1-UD* (SLY2559) cells carrying a chromosomal GFP-tag on the right arm of chromosome IV. The percentage of cells showing cohesion (one dot) or loss of cohesion (two dots) were scored for wild-type and *MCD1-UD* cells for n≥3 experiments. The mean values are plotted on the Y-axis, Error bars indicate standard error of the mean (SEM), ns = non-significant.

### Cohesin sumoylation is required for repression of telomere proximal genes

To test the role of cohesin sumoylation in telomere silencing, we performed a widely used telomere silencing assay using a strain having a *URA3* reporter gene inserted near a modified telomere VR (Fig 1A) (37). *URA3* expression can be qualitatively measured in terms of growth on 5-FOA containing plates. 5-FOA is a counter selection marker for *URA3,* therefore growth on 5-FOA containing plates is inversely proportional to *URA3* expression. *MCD1-UD* or that *MCD1-ud* were introduced in a strain having the *URA3* silencing reporter near telomere VR (Fig 1A) and silencing was assayed using 5-FOA. We observed that *MCD1-UD* was highly sensitive to 5-FOA compared to *MCD1-ud* or wild-type cells, suggesting an increase in *URA3* expression and loss of silencing at telomere VR (Fig 1B).

There is a possibility that *MCD1-UD* may de-sumoylate nearby proteins which are not part of the cohesin complex and de-sumoylation of such non targeted proteins could result in loss of silencing at telomeres. To address this possibility, we created a sumoylation-deficient *MCD1* allele (*mcd1-SD*), by mutating eleven lysine residues to arginine. This allele has been shown to be defective in Mcd1 sumoylation (33). Using the same telomere silencing assay, we found that similar to *MCD1-UD*, *mcd1-SD* cells are also sensitive to 5-FOA (Fig 1C), albeit to a lesser extent, probably because this mutant is defective only in Mcd1 sumoylation and not of other cohesin subunits. These results confirm that the loss of telomere silencing in *MCD1-UD* is due to lack of cohesin sumoylation.

To test whether cohesin sumoylation is required for silencing at natural telomeres, we chose to examine the transcript levels of a native subtelomeric gene, *YFR057w*, by reverse transcriptase real time quantitative PCR (RT-qPCR). We observed that *YFR057w* was significantly de-repressed in *MCD1-UD* as compared to *MCD1-ud* (Fig 1D), further confirming that cohesin sumoylation is indeed required for complete silencing of subtelomeric genes. In addition, *mcd1-SD* cells also showed de-repression of *YFR057w* (Fig 1D).

### Binding of silencing regulators to chromatin is unaffected in cohesin sumoylation deficient cells

Loss of silencing observed in non-sumoylatable cohesin mutants could arise from changes in the expression level of proteins involved in establishment and spreading of silencing.

Silencing at telomeres is thought to be dependent on SIR-complex proteins, Sir2, Sir3 and Sir4. To compare the steady state levels of these silencing proteins, we created Sir3-9Myc and Sir4-9Myc by epitope-tagging them at C-terminus, whereas anti Sir2 antibody was used to detect native Sir2 protein. We did not find any reduction in the steady state expression levels of these proteins by western blot analysis using whole cell lysates (Fig S4).

Furthermore, we did not observe any change in the steady state levels of Rap1 (Fig S4), a factor involved in recruitment of Sir proteins to the telomeres.

As there are no changes in the steady state level of Sir proteins, we examined the binding of Sir2 and Sir4 at *YFR057w,* a gene present near telomere VIR whose expression is increased in the cohesin sumoylation defective cells, by chromatin immunoprecipitation (ChIP). Interestingly, we did not observe significant change in the binding of Sir proteins at this site in the sumoylation deficient mutant compared to *MCD1-ud* (Fig 2A). Previously, we have shown that though Sir binding to telomeres was not altered in cohesin ts mutants, cohesin binding was decreased (26). We tested for such a possibility by analysing Mcd1 binding as proxy for the cohesin complex and observed no change in cohesin binding (Fig 2B) suggesting, that sumoylation affects some other aspect of the complex which is required for transcriptional repression near telomeres.

### Sir proteins are competent to establish silencing in cohesin sumoylation defective cells

As the binding of Sir proteins to the telomeric heterochromatin was not reduced in the mutant, we wondered whether these silencing regulators were competent to establish silencing in this mutant. To test this, we carried out genetic interaction analysis with *RIF1* and *SIR3*. Sir3 overexpression can extend the zone of silencing at telomeres, as it can bind beyond the SIR-complex binding sites independent of other Sir proteins (38,39). Sir3 overexpression also leads to gain of silencing at telomeres (38,39). Consistently, we observed increased silencing of the *URA3* reporter upon overexpression of Sir3 (Fig S5A). Interestingly, Sir3 overexpression in *MCD1-UD* led to complete repression of *URA3* to a level equivalent to the wild-type or *MCD1-ud* (Fig S5A), suggesting Sir proteins are competent to establish silencing and their overexpression supresses the silencing defect of the sumoylation defective cohesin mutant, at least at the modified telomere VR.

Rif1 is a Rap1 interaction factor (40) that competes with Sir4 for binding at the telomeres. In a *rif1* deletion mutant, binding of Sir proteins to the telomeres increases resulting in gain of silencing (41,42). Deletion of *RIF1* resulted in significant enhancement of silencing of the *URA3* reporter in *MCD1-UD rif1* double mutant relative to *MCD1-UD* (Fig S5B). Together, these observations indicate that Sir proteins retain their silencing function in *MCD1-UD* and the transcriptional de-repression in this mutant is not due to a defect in silencing function of the Sir proteins.

### Cohesin sumoylation contributes to subtelomeric gene silencing independent of Sir proteins

Since we did not observe any difference in the association of Sir proteins with the de-repressed subtelomeric gene, we wondered whether the role of cohesin sumoylation in subtelomere silencing could be Sir independent. To test this possibility, we compared the transcript levels of *YFR057w* in *MCD1-UD sir2* double mutant with *MCD1-UD*, *MCD1-ud* (WT) and *MCD1-ud sir2*. We found that *YFR057w* transcripts were more abundant in double mutants relative to either of the single mutant or WT cells (Fig 2C), suggesting that cohesin sumoylation contributes independently of Sir proteins for complete silencing of telomere proximal gene.

### Telomere length maintenance defect in cohesin sumoylation defective cells

Telomeres are repetitive regions at the ends of the chromosomes and their length must be maintained for genome stability. The cell uses various mechanisms to maintain telomere length homeostasis. Telomere length also has a causal relationship with silencing as reduction in telomere length leads to decrease in Rap1 binding site and hence reduced SIR binding and derepression of subtelomeric genes. To test whether derepression of subtelomeric genes is a result of decrease in telomere length, we examined telomere length in *MCD1-UD* using a Y’element specific probe (Fig S6A). Interestingly, we observed an increase in telomere length in *MCD1-UD* compared to wild type cells and cells expressing Mcd1 fused to the UD inactivation mutant (*MCD1-ud*) (Fig S6B), suggesting that decrease in expression of telomere proximal genes is not due to telomere shortening in the cohesin sumoylation defective cells. It also indicates that sumoylation of the cohesin complex is important for telomere length maintenance.

### Analysis of sister chromatid cohesion in the cohesin sumoylation defective cells

Cohesin has an established role in sister chromatid cohesion (23,43), where it tethers two sister DNA molecules to bring about cohesion. We examined sister chromatid cohesion at both chromosome arms and telomeres in *MCD1-UD* cells at 23 °C. Cohesion was assayed using strains expressing GFP or CFP tagged Lac-I, bound to tandem *lacO* repeats integrated at *LYS4* on right arm of chromosome IV (44) or near telomere VIR (45) respectively.

Cohesion was measured in nocodazole treated G2/M arrested cells. When chromatids are cohesed, only a single fluorescent dot is visible whereas loss of cohesion leads to appearance of two dots, each belonging to a sister chromatid (Fig 3A). Using this assay we did not observe any significant change in cohesion, either at the telomere or at the arm site (Fig 3B and 3C respectively), indicating that cohesin sumoylation is not required for sister chromatid cohesion, at least at the tested sites. Interestingly, it also suggests that requirement of cohesin for transcriptional repression of subtelomeric genes is independent of its role in sister chromatid cohesion.

### Cohesin sumoylation is required for telomere tethering to the nuclear envelope

The peripheral nuclear space adjacent to the inner nuclear membrane has been described as a sub-compartment of the nucleus that favours silencing of genes localized in this region (46,47). It has also been proposed that telomeres are tethered to nuclear envelope and that tethering plays an important role in silencing (46,47). Moreover, recently we showed that cohesin is required for telomere tethering to the nuclear envelope (26). To test whether sumoylation is required for this function of cohesin we examined the effect of cohesin de-sumoylation on telomere tethering to the nuclear envelope. We used a strain expressing GFP tagged Lac-I bound to tandem repeats of *lacO* integrated near telomere V-R (45) and the nuclear envelope was visualized by GFP tagging of Nup49 (a nuclear envelope protein). The percentage of nuclei having tethered or untethered configuration was enumerated in wild-type and *MCD1-UD* cells grown at 23 °C and arrested in G2/M with nocodazole. Tethering was calculated by measuring the ratio of distance of telomere dot from nuclear envelope (B) to the diameter of the nucleus (A). A B/A value ranging from 0-0.2 were considered as tethered whereas a B/A value between 0.2-0.5 were considered as untethered (Fig 4A top). We found a modest but statistically significant reduction in tethering of telomere VR to the nuclear envelope and a concomitant increase in untethering in *MCD1-UD* cells (from 31% in WT to 46.45% in mutant) (Fig 4A bottom histogram). This finding indicates that cohesin sumoylation contributes to nuclear membrane tethering of telomeres in G2/M cells.

**Figure 4.**
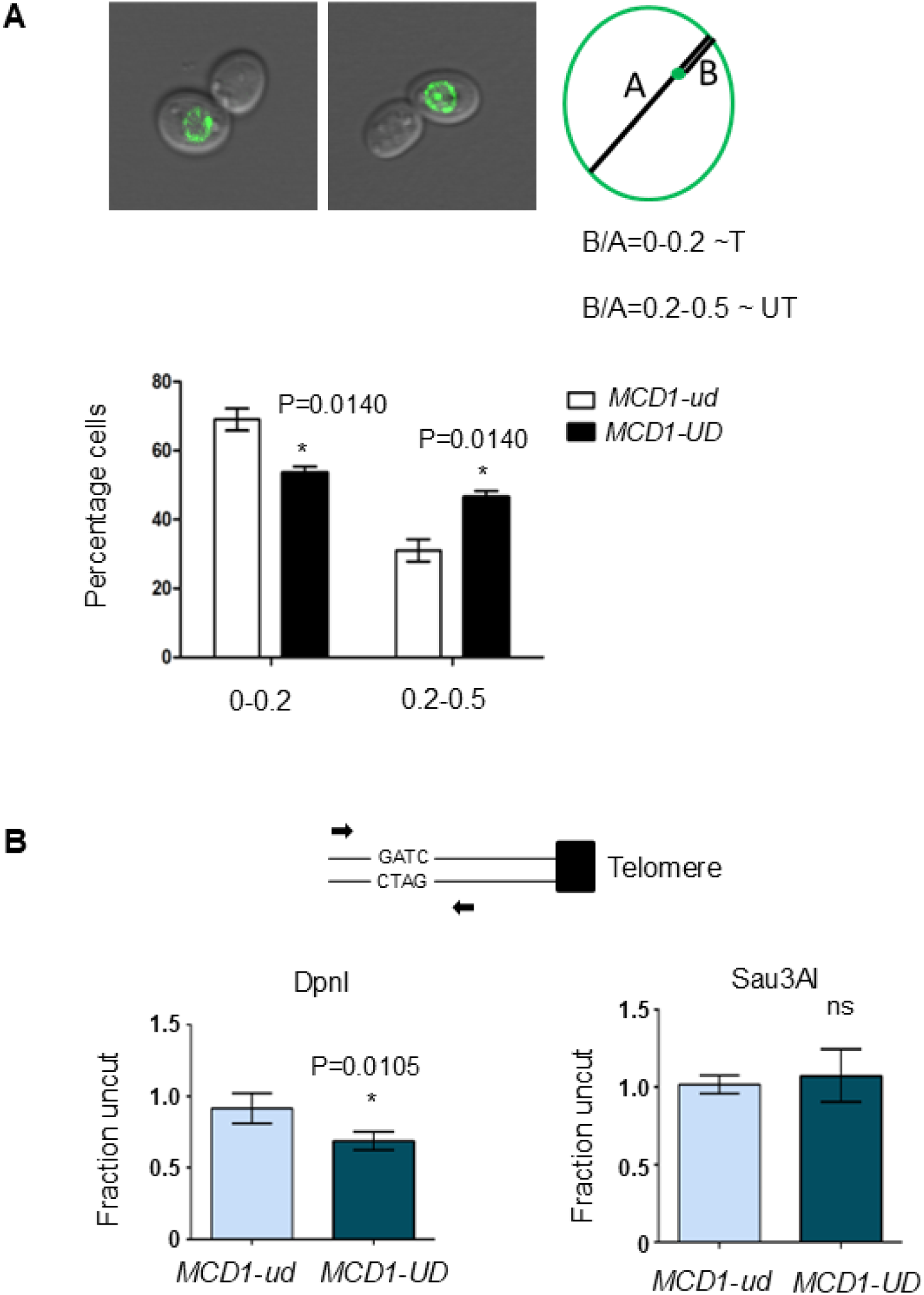
Telomere organization defect in cohesin sumoylation defective cells. (**A**) Tethering of GFP-tagged TEL-VR was measured in nocodazole-arrested *MCD1-ud* (SLY2446) and *MCD1-UD* (SLY2448) strains expressing *NUP49-GFP.* Representative images showing tethered (left) and untethered (right) configurations. A B/A value of 0-0.2 was considered as tethered whereas 0.2-0.5 was considered as untethered (where A is nuclear diameter and B is distance of the telomere dot from the nuclear envelope). The graph showing percentage of cells having tethered versus untethered telomeres is at the bottom. **(B)** Assessment of chromatin accessibility at *YFR057w* by Dam methylase accessibility assay in *MCD1-ud* (SLY2996) and *MCD1-UD* (SLY2998) cells by digestion with DpnI (Left) or Sau3AI (Right) followed by qPCR. The schematic depicts locations of primers (arrows) near TEL-VIR relative to the cutting site. The mean values for n≥3 experiments are plotted on the Y-axis, Error bars indicate standard error of the mean (SEM). The statistical significance was analysed using the two-tailed unpaired t-test where * indicates P≤0.05, ns = non-significant.

### Cohesin sumoylation is required for compaction of the telomere proximal region

Earlier studies demonstrated that telomeres have a special compact organization which is refractory to DNA modifying enzymes (48). Recently we showed that both Sir2 and cohesin are required for the formation of this specialized structure (26). To understand the regulation of this function of cohesin we compared chromatin compaction in *MCD1-ud* and *MCD1-UD* cells by quantifying Dam-methylase (an enzyme that methylates adenine within a GATC sequence) accessibility of sub-telomeric chromatin near Tel-VIR (48). Sub-telomeric chromatin in the sumoylation defective mutant appeared to be more accessible as it was more susceptible to cleavage by Dpn1, indicating a relatively open organization. The same region was equally sensitive to Sau3A1, a methylation insensitive restriction-endonuclease (Fig. 4B). These findings indicate that subtelomeric chromatin in *MCD1-UD* assumes a more open conformation relative to *MCD1-ud* cells. Interestingly, unlike *mcd1-1* non-subtelomeric region (*YFL015C*) remained equally accessible irrespective of sumoylation status of cohesin (Fig. S7).

## Discussion

Cohesin is known to undergo various posttranslational modifications such as acetylation, phosphorylation and sumoylation and these modifications modulate its activity (49–51). Cohesin plays many other important roles beyond its central role in sister chromatid cohesion. Cohesin has a well-established role in regulation of gene expression, and many developmental defects associated with mutations in the cohesin network proteins are believed to be a result of altered gene expression. Surprisingly, no study has directly evaluated the role of posttranslational modifications of cohesin in the regulation of gene expression.

Here, we have investigated the role of cohesin sumoylation in subtelomeric gene silencing. Using *MCD1-UD*, we have shown that cohesin sumoylation is required for silencing of telomere proximal regions (Fig 1), although the effect of cohesin de-sumoylation is not as strong as observed with temperature sensitive mutants of the cohesin complex (26). In contrast to Almedawar *et al*., we found that this fusion construct supports viability and sister chromatid cohesion, at least at the tested loci. These differences could arise due to different strain backgrounds used in these studies. Moreover, Almedawar *et al.* found a decrease in sumoylation of Scc3 and Pds5. We have not tested the sumoylation status of these two proteins with our construct. It is possible that sumoylation of these proteins may not have reduced as much in our conditions and their sumoylation could be important for survival.

We have found that SIR protein binding is retained in sumoylation defective cohesin mutant and that cohesin sumoylation is required independently of Sir proteins, for repression of subtelomeric genes (Fig 2B and C). These observations are consistent with our finding using temperature sensitive mutants of the cohesin complex (26). We also show that cohesin sumoylation is required for telomere organization as sumoylation deficient cohesin mutant has long telomeres (Fig. S6) and is defective in telomere anchoring to the nuclear envelope and telomere compaction (Fig 4A and B).

To understand the role of sister chromatid cohesion in transcriptional silencing of telomere proximal region we analysed cohesion at telomere and an arm region. Interestingly, we observed that cohesion was intact at both the loci. It has been difficult to separate sister chromatid cohesion from other function of cohesin as most of the available mutants are defective in cohesion as well. Importantly, our observation suggest that cohesion and transcriptional silencing are separable function of cohesin. Moreover, no loss of cohesion or change in cohesin binding in sumoylation defective cohesin mutant suggest that cohesin uses a different mechanism for chromatin compaction and sister chromatid cohesion, at least at telomeres.

Based on these observations we propose that sumoylation is required for cohesin’s function in repression of telomere proximal genes independent of its role in cohesion and also independently of the SIR mediated silencing pathway, possibly via its role in telomere organization.

## MATERIALS AND METHODS

For RT-qPCR, RNA was isolated using Qiagen RNeasy minikit and qPCR of cDNA was carried out on a Bio-rad IQ5 real time PCR machine using Bio-rad iTaq universal SYBR green supermix. ChIP was performed as described earlier (10). Confocal microscopy was performed using Zeiss LSM880 (Airyscan) confocal microscope, and were analyzed using ZEN 2.1 (black) software. Detailed experimental procedures are provided in SI (Supporting Information) Materials and Methods.

## ACKNOWLEDGEMENTS

We thank Yves Barral, Susan Gasser, Vincent Guacci, Rohinton Kamakaka, Michael Knop and Mahendrawada Lakshmi for yeast strains and plasmids used in this work. This work was supported by grants from DBT, SERB and CSIR to S. Laloraya and the DBT-IISc partnership program, whose durations are now over. D. Kothiwal was supported by a DBT Senior Research Fellowship. Technical support from the DBT funded confocal microscopy facility of the Division of Biological Sciences, I.I.Sc., is acknowledged. Support for equipment in the Department of Biochemistry is provided by grants from DST-FIST, UGC and I.I.Sc. We thank members of our lab for discussions.

## Author Contributions

D.K. and S.L. designed research; D.K. and S. J. performed research; D.K. contributed new reagents/analytic tools; D.K. and S.L. analyzed data; D.K. and S.L. wrote the paper; and S.L. supervised the work.

