## Supplementary Information for "Cohesin sumoylation is required for repression of subtelomeric gene expression in *Saccharomyces cerevisiae*"

**This PDF file includes:**

- Supporting text
- Figures S1 to S7
- Tables S1 to S3
- SI References

**Other supporting materials for this manuscript include the following:**

Not applicable

### SUPPORTING INFORMATION

#### Materials and methods

##### Media and Reagents

Cells were grown in YPD medium or minimal medium as indicated. We purchased G418 from Sigma, hygromycin from US biological, nourseothricin from Jena bioscience; 12CA5 anti-HA, 9E10 anti-Myc and anti-GFP antibodies from Roche Applied Science, anti-SIR2 antibody (y-80) from Santa-Cruz and HRP-conjugated Anti-mouse and anti-rabbit secondary antibodies from Bangalore Genei.

##### Yeast Strains and Plasmids

The yeast strains used or generated in this study are listed in Table S1. Oligonucleotides used for construction of strains and plasmids were synthesized from Sigma-Aldrich; sequences of these oligonucleotides are provided in Table S2.

##### Integration of *MCD1-UD* or *MCD1-ud* at *MCD1* genomic locus

To integrate *MCD1-UD*, integration cassette (*MCD1-UD*:*LEU2*) was created in three steps. In the first step *MCD1* downstream sequence (sequence after stop codon) was cloned into HincII digested pUC19 to generate pDK16. In second step BglII fragment of pSL617 (*sir2Δ::LEU2*) was cloned into BamHI site of pDK16 to give rise pDK17. In final step *MCD1-UD* was PCR amplified from pDK6 using SLO505 and SLO508 and cloned into SmaI site of pDK17 and named as pDK18. To integrate *MCD1-UD*, HindIII digested fragment of pDK18 was transformed in different yeast strains. Integrations were confirmed by colony PCR using LEU L2B and SLO613. Upon correct integration it gives a band of 610bp.

*MCD1-ud* integration cassette was generated in a similar way, except that *MCD1-ud* was created by site directed mutagenesis using pDK6 as a template and cloned into SmaI site of pDK17, integration and confirmation strategy were also same.

**Integration of *mcd1 SD* or *MCD1*-6HA at *MCD1* genomic locus**

To integrate *mcd1 SD (mcd1 11KR)*, integration cassette (*mcd1 SD-6HA:hphNT1 kanMX6*, pDK28) was created in two steps. In the first step *mcd1 SD-6HA:hphNT1* was created by site directed mutagenesis using *MCD1-6HA:hphNT1* as a template and cloned into SmaI site of pUC19 to generate pDK25. In the second step *kanMX6* containing fragment of pfa6kanMX6 was cloned into BstZI7I site of pDK25 to generate pDK28. To integrate *mcd1 SD*, PsiI digested fragment of pDK28 was transformed in yeast strain ROY783. Integrations were confirmed by colony PCR using SLO791 and SLO613. Upon correct integration it gives a band of ~450bp.

*MCD1-6HA:hphNT1 kanMX6* (pDK33) cassette was created in a similar way. Integration and confirmation strategy were also the same.

**Creation of *RIF1* deletion strain**

*RIF1* deletion cassette was amplified from SLY859 (*rif1Δ::KanMX4*) using SLO882 and SLO883. Yeast cells were transformed with ethanol precipitated PCR product, transformants were selected on G418 plate and deletion was confirmed by colony PCR using SLO441 and SLO884 which gives a band of ~400bp.

**Construction of HF-Smt3 strains**

The plasmid bearing *HF-SMT3* (pTAGRY310) was transformed into W303-1a and genomic copy of *SMT3* was knocked out by PCR mediated one step gene disruption using primers SLO816/SLO817 and template as pYM16. Genomic disruption of *SMT3* was confirmed by colony PCR using primers SLO818/SLO534 which should give 620bp upon disruption.

**Construction of *SIR2* deletion strains**

*SIR2* was deleted using PCR mediated one step gene disruption approach (1). To delete *SIR2* a primer pair SLO679/SLO680 was used with pYM22 as template. Genomic disruption of *SIR2* was confirmed by PCR using oligo pair SLO791/SLO024, which gives a band of ~600bp upon *SIR2* deletion.

**Creation of *SIR3*, *SIR4*, and *RAP1* tagged strains**

All the proteins were epitope tagged at C-terminus using PCR mediated one step tagging approach (1)

*SIR3*: *SIR3* was tagged with 9Myc using primer pair SLO876/877 with pYM20 as template.

*SIR4*: 9Myc Tagging was performed using pYM20 as template with primer pair SLO871/872.

*RAP1*: GFP tagging of *RAP1* was done using pYM27 as template with primers SLO907 and SLO908.

**Spot assay**

To assay growth and drug sensitivity, yeast cells were grown in YAPD until mid-log phase, OD of 0.8-1.0. Cultures were serially diluted ten-fold starting from $10^{-1}$ and spotted on YAPD plates with and without drug (HU, MMS). YAPD plates were kept at various temperatures (23, 30 and 37°C). MMS (0.01%) and HU (100mM) plates were maintained at 23ºc. Plates were incubated for 3-5 days and images were captured at regular interval.

**Preparation of whole cell extracts and immunoblot analysis**

Cells were grown to mid-log phase (OD ~0.8) and protein extracts were prepared by lysing the cells in TCA (trichloro acetic acid) using glass beads. Samples were resolved on SDS-PAGE gel. The resolved proteins were transferred onto nitrocellulose membrane using Bio-Rad semi dry apparatus (39mM glycine, 48mM Tris, and 15% methanol). Proteins were detected using indicated primary antibodies and HRP conjugated secondary antibody. Blots were developed using Perkin-Elmer chemiluminescence reagent.

**Pull-down assay to detect sumoylation**

Pull-down was performed as described previously (2), with minor modifications. To detect His-FLAG tagged SUMO conjugates from yeast cells 200 ml cultures were harvested and disrupted by glass beads in 1 ml Buffer A (8M Urea, 100mM NaH2PO4, 10mM Tris HCl, 0.05% Tween pH 8), and extracts were clarified by centrifugation. Protein concentration was determined using a Bradford Assay, and ~12-15mg of protein was added to 50μl of 50:50 slurry of Ni-NTA beads (prewashed in Buffer A). Imidazole was added to a final concentration of 20mM, to reduce non-specific binding. Proteins were bound for 3 hours at 4°C on a rotating platform. Protein bound beads were washed twice with Buffer A containing 2mM imidazole, followed by 2 washes in Buffer B (8M Urea, 100mM NaH2PO4, 10mM Tris HCl, 0.05% Tween pH 6.3). Bound proteins were eluted off the beads using 30μl of 2X laemili buffer supplemented with 4% β-mercaptoethanol and 200mM EDTA and samples were subjected to SDS-PAGE followed by western blotting.

**Telomere silencing assay**

Overnight grown log phase cultures were normalized to 1 $OD_{600}$ and serially diluted to $OD_{600}$ 0.5, 0.1, 0.01, and 0.001, and spotted on SC and SC 5-FOA (1mg/ml or 0.1%) plates. Plates were incubated at 23 $^{0}$C and images were captured at regular intervals.

**RT-qPCR**

Total RNA was isolated from exponentially growing yeast cells using Qiagen RNeasy minikit. cDNA was prepared from 2ug of DNase treated RNA and qPCR was carried out on a Bio-rad IQ5 real time PCR machine using Bio-rad iTaq universal SYBR green supermix. The primers used for qPCR are listed in table S3. The expression level of test genes in all cases was normalized with respect to *ACT1*. The mean values for $n\geq3$ experiments are plotted on the Y-axis. Error bars represent the standard error of the mean (SEM). The unpaired two-tailed Student's t-test was used to evaluate the statistical significance of the observed differences in expression. * indicates $P\leq0.05$, ** indicates $P\leq0.01$ and *** denotes $P\leq0.001$.

**ChIP-qPCR**

ChIP was performed as described before (3) using asynchronous cultures. qPCR of ChIP samples was done in a Bio-rad IQ5 real time PCR machine using Bio-rad iTaq universal SYBR green supermix. Calculations were done using ΔΔCt methods where Ct values at test locus were normalized with respect to total and a control locus (As indicated in Figure legend) (4). The primers used for qPCR are listed in Table S3. The fold enrichment is relative to the *PHO5* in all ChIP-qPCR experiments. The mean values for $n\geq3$ experiments are plotted on the Y-axis. Error bars represent the standard error of the mean (SEM). For simplicity, the wild type values were normalized to 1.

**Southern Blot**

Genomic DNA for Southern blot analysis was isolated from exponentially growing yeast cell cultures by glass bead lysis (5). The DNA pellet obtained was resuspended in 40-50 μl of sterile water. 6.0-8.0 μg of the Xho1 digested genomic DNA was electrophoresed on a 1% agarose gel in 1X TAE (40mM Tris 20mM acetate, 1 mM EDTA) buffer and then transferred onto Hybond-N nylon membranes (GE Healthcare) overnight at room temperature using 10X SSC (1.5 M NaCl, 0.15 M Na-citrate) as the transfer buffer. The blot was uv-crosslinked using Amersham UVC 500 crosslinker for 5 minutes. To examine telomere lengths, the membrane was probed with a $^{32}$P radiolabelled probe specific for the Y' element, obtained from pRR48. The radiolabeled probe was generated by random priming using α-$^{32}$P dATP in a reaction containing template DNA (25-50 ng), 0.5mM dNTPs, Klenow polymerase buffer, random primer and Klenow Polymerase. Signal intensities were obtained by detecting photo-stimulated luminescence counts in a GE Typhoon FLA9000 phosphorimager.

**Microscopy**

For the cohesion assay, exponentially growing cultures were supplemented with 1% DMSO, for 30 minutes and nocodazole (SIGMA) was added to a final concentration of 15μg/ml. Metaphase-arrested cells carrying GFP or CFP chromosome tags were analyzed by confocal microscopy, where 3 μl (1 mg/ml) DAPI was added for DNA staining 20 minutes prior to harvesting. Cells were immediately fixed with 4% paraformaldehyde and imaging was performed using fixed cells. Experiment was repeated three times and each time ~100 cells were counted for both Wild-type and mutant.

Similarly for telomere tethering, exponentially growing cultures were supplemented with 1% DMSO, for 30 minutes and nocodazole (SIGMA) added to a final concentration of 15μg/ml. Metaphase-arrested cells carrying *NUP49-GFP* and GFP chromosome tag were analyzed by confocal microscopy. Cells were immediately fixed with 4% paraformaldehyde and imaging was performed using fixed cells. Experiment was repeated three times with two individual biological isolates. Each time ~50 cells were counted for all the samples.

In both cohesion and tethering experiments images were captured, using LSM880 (Airyscan) confocal system from Zeiss, and were processed using ZEN BLACK 2.1 software.

**Telomere accessibility assay**

Telomere accessibility assay was performed as before (*). Yeast strains expressing the *E. coli dam* methylase (that methylates adenine within GATC) were grown to log phase and genomic DNA was extracted. Samples were digested overnight at 37°C with DpnI or Sau3AI (methylation sensitive or insensitive restriction endonucleases that recognize GATC) and analyzed by quantitative real-time PCR. Primers were designed such that the telomere VIR proximal PCR-fragment generated includes the DpnI/Sau3AI restriction site (~1.2 kb from the telomere). The fraction of uncut DNA relative to wild-type was measured by qPCR and normalized to an uncut DNA fragment at *ACT1*. Normalization was performed using a primer set (*ACT1-dam*-F/*ACT1-dam*-R**)** amplifying an uncut sequence at the *ACT1* gene. The unpaired two-tailed Student's t-test was used to evaluate the statistical significance of the observed differences in accessibility. * indicates P≤0.05, ** indicates P≤0.01 and *** denotes P≤0.001.

**SI Figures S1 to S7**

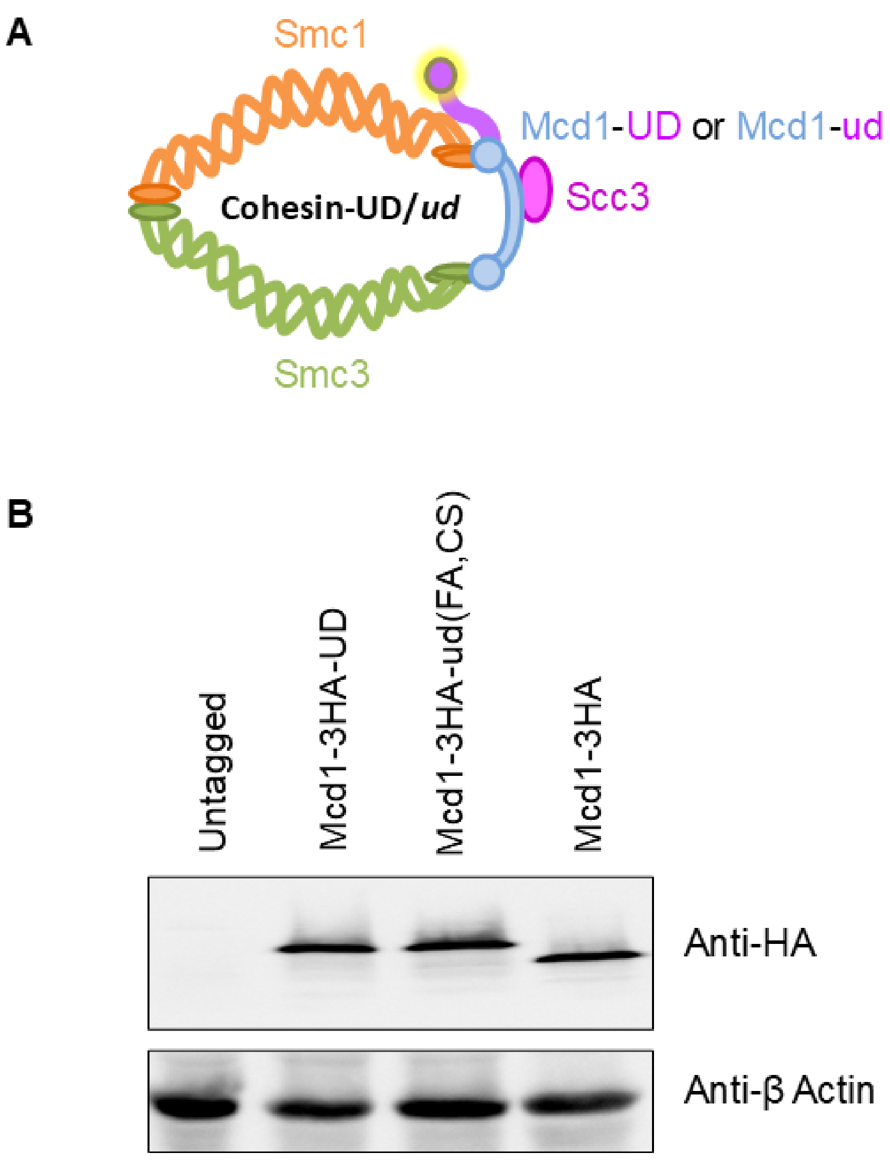


**Figure S1. Construction of Mcd1-3HA-UD fusion. (A)** Schematic showing fusion of Ulp1 domain, UD (purple) to the kleisin component of cohesin complex, Mcd1 (blue). **(B)** Western blot showing equivalent expression levels of Mcd1-3HA (SLY1807), Mcd1-3HA-

UD (SLY1820) and Mcd1-3HA-ud(FA,CS) (SLY1823). β Actin was used as a loading control.

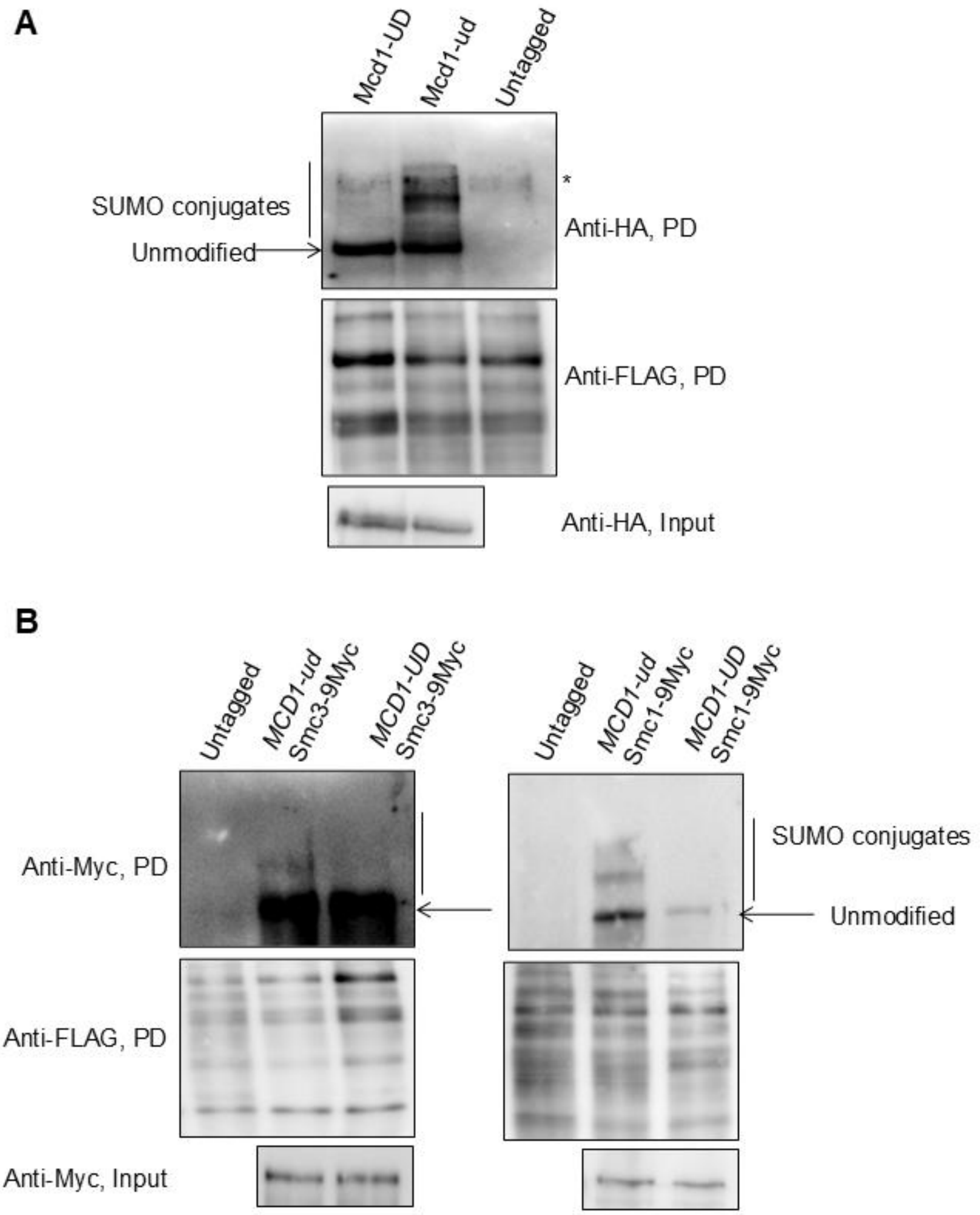


**Figure S2. Fusion of ULP domain (UD) can de-sumoylate the cohesin complex.** Analysis of sumoylation of different subunits of the cohesin complex. Pull-down assays (PD) were carried out using a strain (SLY2480) expressing SUMO with 6His-Flag tag at the N-terminus. All the sumoylated proteins in the cell were pulled down using Ni-NTA resin. **(A)** Fusion of active UD (Mcd1-UD) downregulates Mcd1 sumoylation. HF-SUMO strains expressing Mcd1-UD (SLY2905) or Mcd1-ud (SLY2906) were used. After pull-down, Anti-FLAG blot shows the pull-down efficiency and the anti-HA blot detects the sumoylated species. **(B)** Pull down and western blot to detect Smc1-9Myc (SLY2907 and SLY2908) and Smc3-9Myc (SLY2909 and SLY2910) sumoylation in Mcd1-UD or Mcd1-ud backgound. Note that sumoylation of both the proteins decreased in Mcd1-UD.

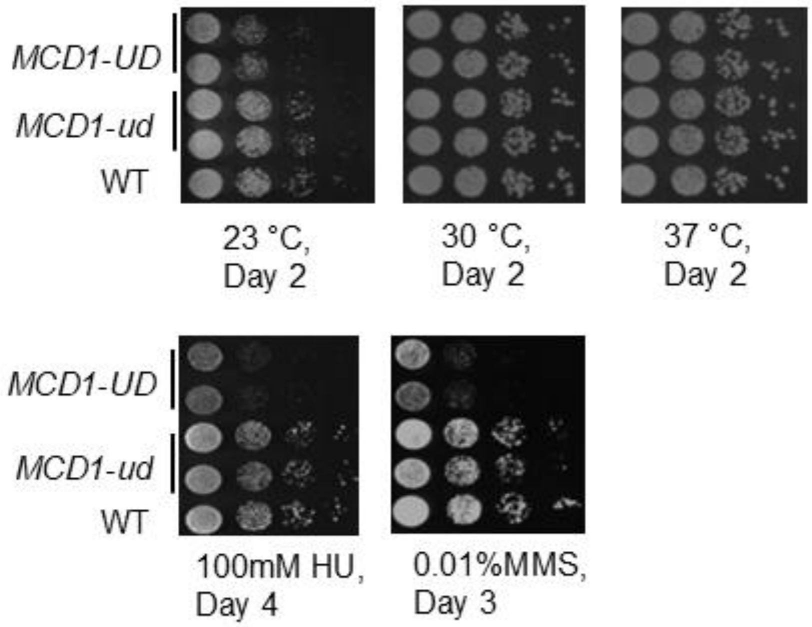


**Figure S3. Characterization of growth defect of *MCD1-UD*.** Exponentially growing cells were serially diluted (10-fold) and spotted on YPD and YPD containing indicated drugs. Images were captured at the indicated times.

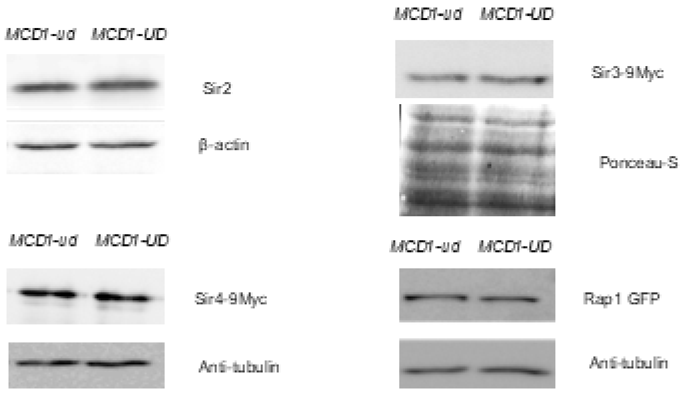


**Figure S4. Study state levels of silencing regulators in not altered in cohesin sumoylation deficient strains**. Western blot showing equivalent levels of Sir2 in *MCD1-ud* (SLY1823) and *MCD1-UD* (SLY1820) strains, Sir3-9Myc in *MCD1-ud* (SLY2395) and *MCD1-UD* (SLY2396) strains, Sir4-9Myc (in SLY2387 and SLY2388 that are *MCD1-ud* and *MCD1-UD* respectively) and Rap1-6HA in *MCD1-ud* (SLY2455) and *MCD1-UD* (SLY2457) strains.

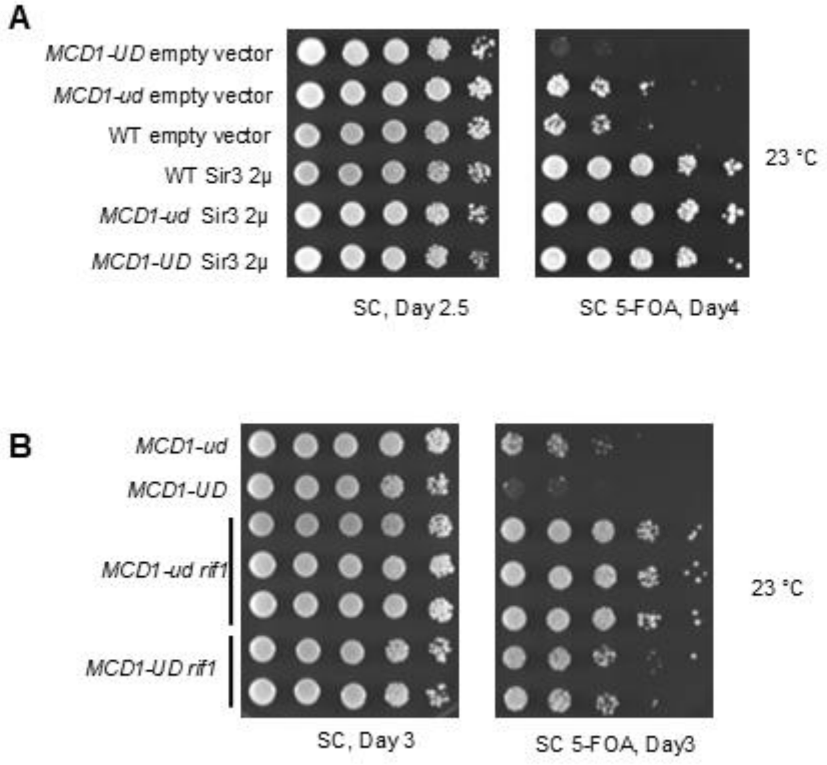


**Figure S5. Sir proteins are silencing competent in *MCD1-UD*.** Genetic analysis of Sir protein function in *MCD1-UD,* using a strain bearing telomere proximal *URA3.* (**A**) Sir3 overexpression suppresses the silencing defect in *MCD1-UD.* TPE determination upon Sir3 overexpression from a 2μ plasmid (pRO146, a kind gift from R. Kamakaka). Overnight grown log phase cultures were serially diluted and spotted onto SC-Trp and SC-Trp + 0.1% 5-FOA plates at 23 °C in wild type (ROY783), *MCD1-ud* (SLY1823) and *MCD1-UD* (SLY1820) strains. (**B**) Suppression of TPE defect by *RIF1* deletion. Dilution spotting for TPE analysis on SC and SC + 0.1% 5-FOA plates at 23 °C in *MCD1-ud* (SLY1823), *MCD1-UD* (SLY1820), *MCD1-ud rif1* (SLY2526), *MCD1-UD rif1* (SLY2528).

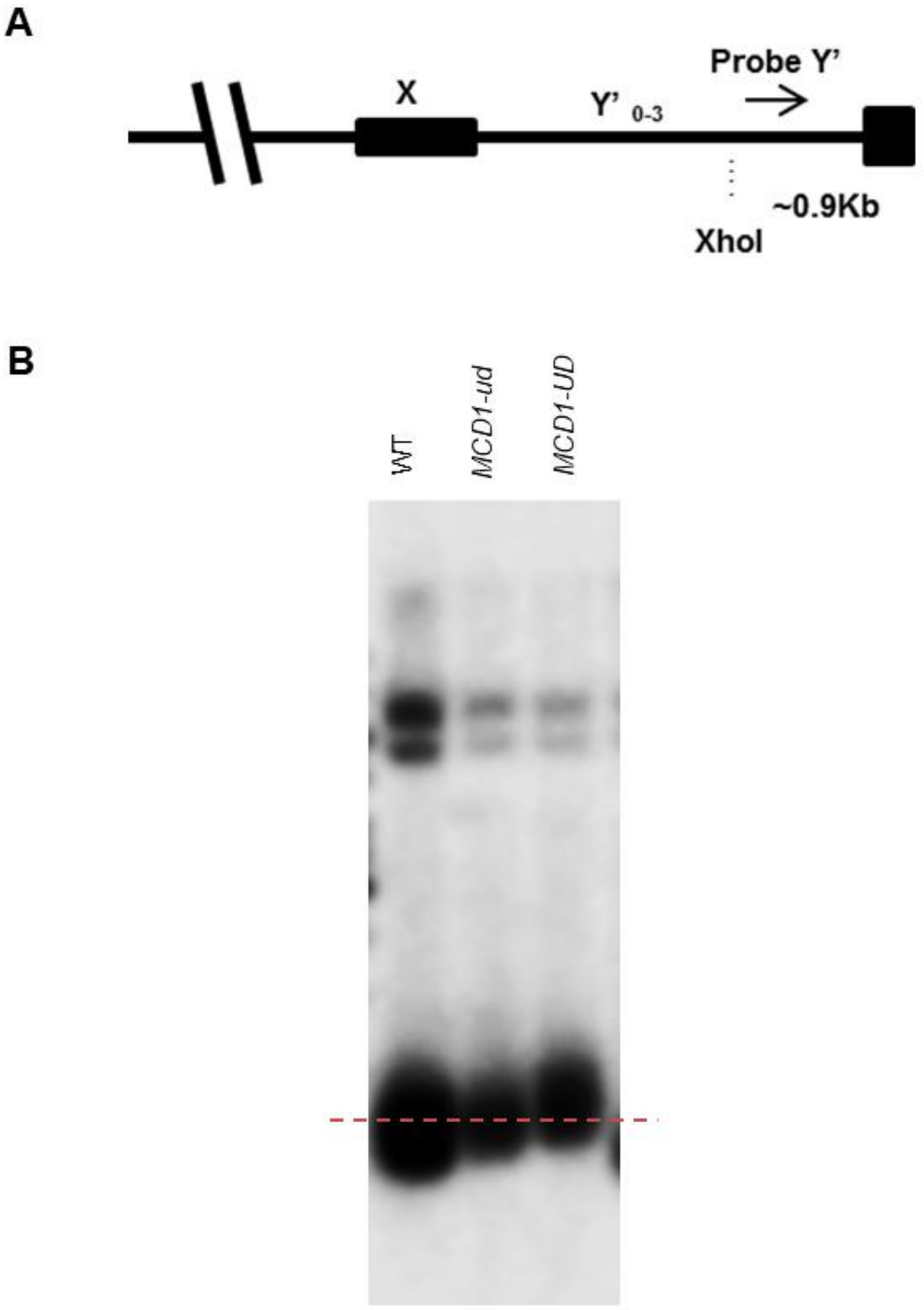


**Figure S6. Telomere length analysis in *MCD1-UD*.** **(A)** Schematic depicting a yeast chromosome end, showing the location of the probe fragment relative to the XhoI restriction site and a Y' subtelomeric element. **(B)** Southern blot analysis for telomere length measurement in WT (SLY1807), *MCD1-ud* (SLY1823) and *MCD1-UD* (SLY1820).

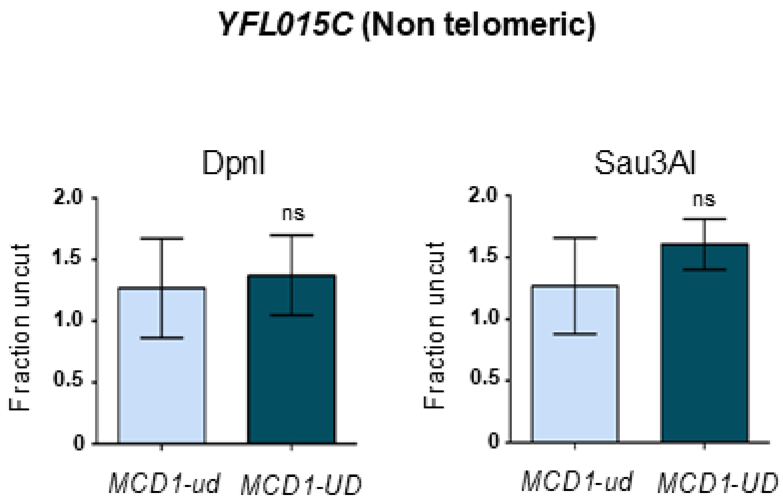


**Figure S7. Assessment of chromatin accessibility at a telomere distal region, *YFL015C*.** Dam methylase accessibility assay was carried out using *MCD1-ud* (SLY2996) and *MCD1-UD* (SLY2998) cells by digestion with DpnI (Left) or Sau3AI (Right) followed by qPCR. The mean values for n≥3 experiments are plotted on the Y-axis, Error bars indicate standard error of the mean (SEM), ns = non-significant.

#### SI Tables S1 to S3

**Table S1**

Yeast Strains used in this study

| Strain | Genotype | Source/Ref. |
|---|---|---|
| ROY 783 | *MATa ADE2 leu2-3,112 his3-11,15 ura3-1 trp1-1 ppr1D::HIS3 URA3-TEL-VR* | Donze D 1999 G&D |
| SLY1820 | ROY 783 *MCD1-3HA-UD:LEU2* | This study |
| SLY1823 | ROY 783 *MCD1-3HA-ud:LEU2* | This study |
| SLY1807 | JS311 *MCD1-3HA:TRP1* | This study |
| SLY2480 | *MATa ade2-1 ura3-1 his3-11,15 trp1-1 leu2-3,112 can1-100 smt3Δ::hphNT1*, pYRTAG310 | Lakshmi Mahendrawada |
| SLY2905 | SLY2480 *MCD1-3HA-UD:LEU2* | This study |
| SLY2906 | SLY2480 *MCD1-3HA-ud:LEU2* | This study |
| SLY2907 | SLY2905 *SMC1-9Myc:natNT2* | This study |
| SLY2908 | SLY2906 *SMC1-9Myc:natNT2* | This study |
| SLY2909 | SLY2905 *SMC3-9Myc:natNT2* | This study |
| SLY2910 | SLY2906 *SMC3-9Myc:natNT2* | This study |
| SLY2743 | ROY783 *MCD1-6HA:hphNT1 KanMX6* | This study |
| SLY2745 | ROY783 *mcd1 SD-6HA:hphNT1 KanMX6* | This study |
| SLY2387 | SLY1823 *SIR4-9Myc:hphNT1* | This study |
| SLY2388 | SLY1820 *SIR4-9Myc:hphNT1* | This study |
| SLY2395 | SLY1820 *SIR3-9Myc:hphNT1* | This study |
| SLY2396 | SLY1823 *SIR3-9Myc:hphNT1* | This study |
| SLY2450 | W303-1a *RAP1-GFP:hphNT1* | This study |
| SLY2455 | W303-1a *MCD1-ud:LEU2 RAP1-GFP:hphNT1* | This study |
| SLY2457 | W303-1a *MCD1-UD:LEU2 RAP1-GFP:hphNT1* | This study |
| SLY2520 | ROY783 *rif1Δ::KanMX4* | This study |
| SLY2526 | SLY1823 *rif1Δ::KanMX4* | This study |
| SLY2528 | SLY1820 *rif1Δ::KanMX4* | This study |
| SLY2911 | SLY1823 *sir2Δ::TRP1* | This study |
| SLY2912 | SLY1820 *sir2Δ::TRP1* | This study |

| | | |
|---|---|---|
| GA2201 | *MATa ade2-1 can1-100 his3-11,-15 leu2-3,-112 trp1-1 ura3-1 ade2-1::HIS3p-CFP-lacI-URA3p-tetR-YFP-ADE2 TELVI-L::tetO-LEU2, TELVI-R::lacO-TRP1* | Kerstin Bystricky 2005 JCB |
| SLY2446 | GA2201 *Mcd1-3HA-ud:KanMX4* | This study |
| SLY2448 | GA2201 *Mcd1-3HA-UD:KanMX4* | This study |
| yYB3476 | *MATa ura3-52 his3Δ200 leu2 lys2-801 ade2-101 trp1Δ63 trp1::TetO:TRP1 lys4::LacO:LEU2 his3::LacR-GFP:HIS3 TetR-mRFP* | Neurohr G 2011 Science |
| SLY2549 | yYB3476 *Mcd1-3HA-ud:KanMX4* | This study |
| SLY2551 | yYB3476 *Mcd1-3HA-UD:KanMX4* | This study |
| GA2198 | *MATa ade2-1 can1-100 his3-11,-15 leu2-3,-112 trp1-1 ura3-1 his3-11,-15::HISp-GFP-LacI-HIS3, nup49::NUP49-GFP TELV-R::lacO:TRP1* | Kerstin Bystricky 2005 JCB |
| SLY2557 | GA2198 *Mcd1-3HA-ud:LEU2* | This study |
| SLY2559 | GA2198 *Mcd1-3HA-UD:LEU2* | This study |
| SLY2996 | *MAT*a *leu2-3,112 trp1-1 can1-100 ura3-1 ade2-1 his3-11,15 Mcd1-3HA-ud:LEU2* $dam^{+}$ *hphNT1* | This study |
| SLY2998 | *MAT*a *leu2-3,112 trp1-1 can1-100 ura3-1 ade2-1 his3-11,15 Mcd1-3HA-UD:LEU2* $dam^{+}$ *hphNT1* | This study |

**Table S2**

Oligonucleotides used in this study

| | |
|---|---|
| SLO-024 | TCTATACTGATACCACGCCT |
| SLO-441 | CGCTATACTGCTGTCGATTC |
| SLO-534 | GTATCTAGCAGCAGAACCGG |
| SLO-603 | GAAATATTAAAATAGACGCCAAACCTGCACTATTTGAAAGGTTTATC AATGCTCGTACGCTGCAGGTCGAC |
| SLO-604 | GAAATATTAAAATAGACGCCAAACCTGCACTATTTGAAAGGTTTATC AATGCTCGTACGCTGCAGGTCGAC |
| SLO-613 | CTCTGGCTGCTAATGTACC |
| SLO-679 | CTTCGGTAGACACATTCAAACCATTTTTCCCTCATCGGCACATTAAAG CTGGATGCGTACGCTGCAGGTCGAC |
| SLO-680 | TATGTAAATTGATATTAATTTGGCACTTTTAAATTATTAAATTGCCTT CTACTTAATCGATGAATTCGAGCTCG |
| SLO-791 | CGAGCTCGAATTCATCGAT |
| SLO-816 | GGACAGAAGGACCCAGTTCAGTTCTAGTTTTACAAATAAATACACGA GCGCGTACGCTGCAGGTCGAC |
| SLO-817 | TGGGGGGAAGGGAGAGGTTTGTTGGCGTTTTTTAGGCATTGTTAAGA GTCATCGATGAATTCGAGCTCG |
| SLO-818 | GAGTTGCCGTGCCTTTCC |
| SLO-871 | ATTAACAAATTGATGGAAAAAGATTTTCAAGTGAATAAGGAGATAA AACCGTATCGTACGCTGCAGGTCGAC |
| SLO-872 | CAAAGAAAAACAGGGTACACTTCGTTACTGGTCTTTTGTAGAATGAT AAAAAGTCAATCGATGAATTCGAGCTCG |
| SLO-876 | CTGCATGTGTACATAGGCATATCTATGGCGGAAGTGAAAATGAATGT TGGTGGTCAATCGATGAATTCGAGCTCG |
| SLO-877 | GAAATAAATTACGCCTTTTCGATGGATGAAGAATTCAAAA ATATGGACTGCATTCGTACGCTGCAGGTCGAC |
| SLO-882 | GCATTATAATCGCGAAAAAG |
| SLO-883 | ATTATCCTAATTCCCAACTG |

| | |
|---|---|
| SLO-884 | CGGTAGCATTTCCATCATA |
| SLO-907 | GGAACTGGTAGAATGGAAATGAGGAAAAGATTTTTTGAGAAGGACC TGTTACGTACGCTGCAGGTCGAC |
| SLO-908 | AAATAAAGGAGTAAAATAAGTTAAACAATGATGTTACTTAATTCAAT TACTCAATCGATGAATTCGAGCTCG |
| SL-LEU-L2B | CCTTGCGTTTCAGCTTCCACTA |

**Table S3**

qPCR primers used in this study

| Primer | Sequence | |
|---|---|---|
| SLO-853 | CTAGTGTCTATAGTAAGTGCTCGG | *YFR057w* FP |
| SLO-854 | GGTATATTGCCACGCAAAGAAAGG | *YFR057w* RP |
| SLO-861 | GCCGGTATTGCCGCTGCC | *PAU3* FP |
| SLO-862 | CCTTAGATAGAGCACTGGAGATG | *PAU3* RP |
| SLO-867 | GGAAACGTAGAAGGCTGGAACGTT | *ACT1* FP |
| SLO-868 | ACAACGAATTGAGAGTTGCCCCAG | *ACT1* RP |
| SLO-859 | GAATCGATACAACCTTGGCACTC | *PHO5* FP |
| SLO-860 | GGTAATCTCGAATTTGCTTGCTC | *PHO5* RP |
| SLO-1025 | GCTGAGATAAGTAATATCGTTGATGAA | TEL6R-*dam*-F |
| SLO-1026 | TCAAACAAGTAGGAATGCGAAAG | TEL6R-*dam*-R |
| SLO-1027 | AGGTTGCTGCTTTGGTTA | *ACT1*-*dam*-F |
| SLO-1028 | CGTAGGAGTCTTTTTGACCC | *ACT1*-*dam*-R |
| SLO-1080 | TCAACACGTCGTCATTGC | NT-*dam*-FP |
| SLO-1081 | CGCAAATCCAAGTGAAAATC | NT-*dam*-RP |
